# GGTyper: genotyping complex structural variants using short-read sequencing data

**DOI:** 10.1101/2024.03.15.585230

**Authors:** Tim Mirus, Robert Lohmayer, Bjarni V. Halldórsson, Birte Kehr

**Affiliations:** Leibniz Insitute for Immunotherapy, 93053 Regensburg, Germany; deCODE genetics/Amgen Inc., Reykjavík, Iceland; University of Regensburg, 93053 Regensburg, Germany

## Abstract

**Motivation:** Complex structural variants are genomic rearrangements that involve multiple segments of DNA. They contribute to human diversity and have been shown to cause Mendelian disease. Nevertheless, our abilities to analyse complex structural variants are very limited. As opposed to deletions and other canonical types of structural variants (SVs), there are no established tools that have explicitly been designed for analysing complex SVs.

**Results:** Here, we describe a new computational approach that we specifically designed for genotyping complex SVs in short-read sequenced genomes. Given a variant description, our approach computes genotype-specific probability distributions for observing aligned read pairs with a wide range of properties. Subsequently, these distributions can be used to efficiently determine the most likely genotype for any set of aligned read pairs observed in a sequenced genome. In addition, we use these distributions to compute a genotyping difficulty for a given variant, which predicts the amount of data needed to achieve a reliable call. Careful evaluation confirms that our approach outperforms other genotypers by making reliable genotype predictions across both simulated and real data. On up to 7829 human genomes, we achieve high concordance with population-genetic assumptions and expected inheritance patterns. On simulated data, we show that precision correlates well with our prediction of genotyping difficulty. This together with low memory and time requirements makes our approach well-suited for application in biomedical studies involving small to very large numbers of short-read sequenced genomes.

**Availability:** Source code is available at https://github.com/kehrlab/Complex-SV-Genotyping.

## 1 Introduction

A major portion of human genetic diversity is due to structural variants (SVs) (Ebert et al., 2021). Only recently, the number of SVs that can be reliably detected per human genome has increased from thousands (Sudmant et al., 2015; Collins et al., 2020; Halldorsson et al., 2022) to tens of thousands (Audano et al., 2019; Beyter et al., 2021) by using long-read sequencing data. This demonstrates that the majority of human SVs have remained undetected in previous studies using short-read data. Nevertheless, recent studies have shown that genotyping of a significant number of the “hidden” variants is possible with short-read data (Aganezov et al., 2022; Liao et al., 2023). In other words, determining a genotype from short-read data when the variant is already known can be feasible even if current short-read approaches fail to detect a variant. To make full use of the massive amount of short-read data that is available today, approaches are needed for efficiently and accurately genotyping known variants, which have been detected in the, as of today, more limited number of long-read sequenced genomes.

SVs are commonly defined as large differences between genomes that encompass at least 50 base pairs of sequence. Unless non-reference sequence is inserted, SVs can be described in terms of breakpoint positions on the reference genome and novel junctions. More precisely, we refer to a position on the reference genome, where the ends of two segments of DNA that are rearranged in a variant connect to each other, as a breakpoint. A novel junction is the connection of two DNA segments that are not adjacent in the reference genome but on a variant allele. Canonical SVs (i.e. deletions, duplications, insertions, inversions, and translocations of a single DNA segment) are associated with fixed numbers of breakpoints and novel junctions (e.g. a deletion has two breakpoints and one novel junction). Complex SVs are variants that involve multiple segments of DNA and therefore cannot be described by any single canonical type. They can be formed by almost arbitrary numbers of breakpoints and novel junctions.

Since their first discovery, it has been uncontroversial that SVs are of great biomedical relevance (Lupski, 1998): Germline SVs affect the risk of many complex diseases (Collins et al., 2020), can be the cause of rare inherited diseases (Lupiáñez et al., 2015), and somatic SVs play a central role in many cancers (Li et al., 2020). Complex SVs have been studied primarily in the context of somatic variation in cancer genomes (Li et al., 2020) and received limited attention in germline genome analysis (Sanchis-Juan et al., 2018). While the number of complex SVs detected in germline genomes is small compared to canonical SVs (e.g. in gnomAD, 5 295 of 433 371 SVs are complex (Collins et al., 2020)), complex germline SVs have been found to be more abundant than previously appreciated (Collins et al., 2017). It follows that complex SVs should be considered when studying the genetic causes of human disease, e.g., in genome-wide association studies.

Stand-alone genotyping approaches are needed for genotyping known variants in large numbers of samples. Many SV detection tools for short-read data, among them some specifically designed for complex SVs (Zhao et al., 2016), do not compute genotypes at all (Layer et al., 2014) or do not separate genotyping from detection (Chen et al., 2015; Zhou et al., 2023). For complex SVs, there is a surprising lack of genotyping approaches that can handle their diverse structures and output a single genotype per variant.

The classic genotyping approach relies on an alignment of the reads to a linear reference genome and leverages paired-end and split-read information for genotyping SVs (Rausch et al., 2012; Chiang et al., 2015; Ebler et al., 2017; Collins et al., 2020). More recently, graph-based approaches for genotyping SVs have emerged. Some of them (Eggertsson et al., 2019; Chen et al., 2019) locally realign reads to the variants’ graph representation in order to determine an allele of origin per read. Others require a read alignment to a pangenome graph as input (Hickey et al., 2020), computed for example with the vg toolkit (Garrison et al., 2018; Sirén et al., 2021). Finally, *k*-mer based genotyping approaches (Sibbesen et al., 2018; Ebler et al., 2022) count haplotype-specific *k*-mers in unaligned read data.

Most genotypers that follow the classic approach or the graph-based approach with realignment can only genotype canonical SVs or individual novel junctions but not entire complex SVs. Paragraph (Chen et al., 2019) is a notable exception but uses a heuristic approach to combine possibly conflicting calls from several novel junctions and has not been benchmarked on complex SVs. Pangenome-based approaches require all variants subject to genotyping to be known and be part of the graph before performing the computationally expensive read alignment. In many application scenarios, especially when variants of interest are rare, this may not be the case. Similarly, *k*-mer based approaches suffer from long running times for genotyping a relatively small number of variants in genomes with already aligned read data and are vulnerable to the smallest breakpoint inaccuracies.

Here, we set out to develop a generic method that uses aligned short-read data to genotype known, breakpoint-resolved, complex SVs which do not contain any non-reference sequence. We approach the problem of genotyping complex SVs as a generalization of the problem of genotyping canonical variants. Our central contribution is the computation of genotype-specific probability distributions for read pairs given the description of a complex SV. These pre-computed distributions allow us to predict the difficulty of distinguishing the genotypes of a given SV from each other. Using a standard maximum-likelihood model, we use the distributions to determine the most likely genotype and a bootstrap certainty for given sets of observed read alignments. We implemented these methods in the tool GGTyper (Generalized GenoTyper), which can genotype both canonical and complex SVs. GGTyper outputs a single genotype per SV independently of the SV’s number of breakpoints and novel junctions. Evaluation on simulated and real data demonstrates consistently high accuracy while keeping time and memory requirements low. This makes GGTyper ideally suited to genotyping known complex SVs in many short read sequenced genomes.

## 2 Algorithm

We present a new genotyping algorithm for biallelic complex SVs given an individual’s aligned short-read sequencing data and a description of the variants’ novel junctions. In addition to computing genotype likelihoods, we propose a bootstrapping-based genotype certainty and describe how our approach allows us to estimate the difficulty of genotyping a given variant.

### 2.1 Genotype likelihoods

The basis of our genotyping algorithm is a maximum-likelihood model that uses the variant structure and the empirical insert-size distribution of the sequencing library for determining which of the three possible genotypes fits the observed alignment data best. Given the set of observed read pairs ℛ, we use Bayes’ theorem and the assumption of independence among all read pairs to define the likelihood of genotype *G*_*i*_, *i* ∈ {0, 1, 2},as

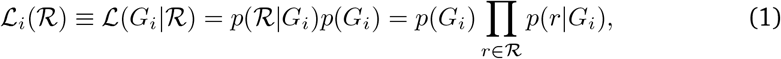

where *p*(*r*|*G*_*i*_) denotes the probability of observing read pair *r* for a given genotype *G*_*i*_. We assume the prior probabilities of the variant *p*(*G*_*i*_) for all three genotypes to be equal unless otherwise specified. Then, the most likely genotype for the observed data is *G*_*k*_ with *k* = argmax_*j*∈{0,1,2}_ ℒ_*j*_(ℛ).

Our central contribution is the derivation of the probabilities *p*(*r*|*G*_*i*_) as presented below. In short, we create allele-specific read-pair profiles by enumerating and weighting all combinations of read pair categories and 5’-differences at all relevant positions along the variant and reference allele. By mixing the allele-specific profiles into genotype-specific profiles followed by normalization, we obtain the probabilities *p*(*r*|*G*_*i*_).

#### 2.1.1 Read pair 5’-differences

We define a read pair’s *5’-difference d* as the (signed) difference between the alignment positions of the 5’ ends of the two aligned reads in the pair. If one read aligns in forward and the other in reverse complement orientation (*FR* and *RF* pairs), we use the position of the reverse read’s 5’-end minus the forward read’s 5’-end (see Fig. S8). By concatenating all chromosomes in a defined order and orientation, we can compute the 5’-difference even if the two reads in a pair align to different chromosomes. In the vicinity of novel junctions, the 5’-difference derived from an alignment can differ from the *insert size s*, which we define as the length of the actual, sequenced DNA fragment.

#### 2.1.2 Read pair categories

We categorize aligned read pairs based on their orientation and location on the reference genome relative to breakpoints ℬ and novel junctions 𝒥. More specifically, we say that a read pair is *split due to novel junction j* ∈𝒥 if the primary alignment to the reference genome of at least one read in the pair is clipped at one of the variant breakpoints that corresponds to the novel junction. As we consider only primary alignments, a read pair can be split at up to four novel junctions. Similarly, we say that a read pair *spans breakpoint b* ∈ ℬ if at least one read in the pair aligns to the reference genome across *b*. For biallelic variants, we define one category per subset *J* ⊆ 𝒥or *B* ⊆ ℬ. In addition, we define one category per orientation *o* ∈ {*FR, FF, RR*} for read pairs neither split due to any junction nor spanning any breakpoint. We chose these categories such that read pairs assigned to a category other than *FR* discriminate between the reference and the variant allele. We ensure that categories are mutually exclusive by assigning read pairs to categories using an order of precedence that is described in the Supplementary Material Sec. 2.4.

#### 2.1.3 Allele-specific read-pair profiles

We define two allele-specific profiles, *ρ*^VAR^ for the variant allele and *ρ*^REF^ for the reference allele. Intuitively, the profiles specify the expected amount of read pairs from the respective allele per category and 5’-difference. In other words, *ρ*^VAR^ and *ρ*^REF^ consist of one 5’-difference histogram of reference-aligned read pairs per read-pair category (Fig. 1C).

**Figure 1.**
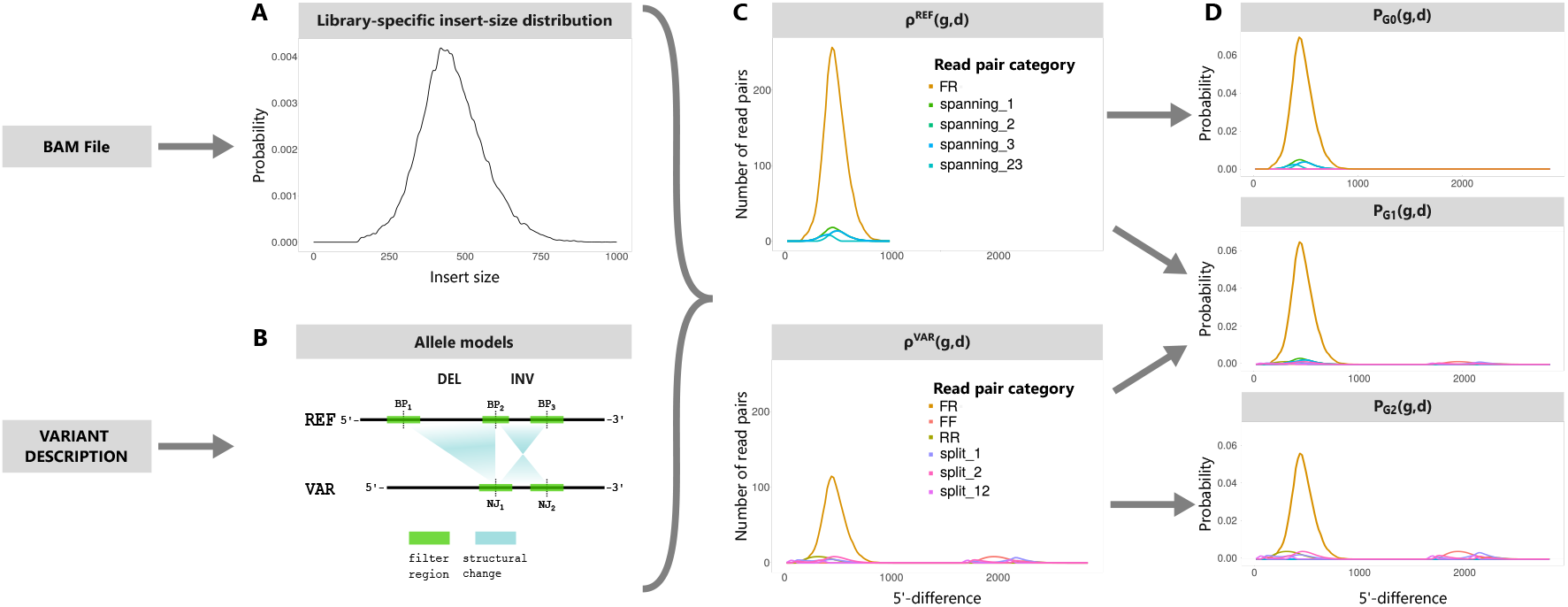
Overview of our approach for calculating read-pair probabilities. We estimate an insert-size distribution from the BAM file (A). We create a reference (REF) and a variant (VAR) allele model from the variant description (B). We calculate allele-specific read-pair profiles from the insert-size distribution and the allele models (C). These profiles contain expected read-pair occurrence frequencies for all possible combinations of read-pair categories and 5’-differences. We obtain genotype profiles by mixing allele-specific read-pair profiles (D). BP, breakpoint. NJ, novel junction. split_xy, read pairs split due to novel junctions x and y. spanning_xy, read pairs spanning breakpoints x and y. FF, forward-forward read pairs. FR, forward-reverse read pairs. RR, reverse-reverse read pairs.

Our idea for creating *ρ*^VAR^ and *ρ*^REF^ is to sample read pairs from regions of interest of either allele (Fig. 1B) according to the empirical insert-size distribution of the sequencing library *P*_Δ_ (Fig. 1A) and considering 5’-differences and categories after mapping to the reference genome. The regions of interest *R*_*ℬ*_ = {[*x*_*b*_, *y*_*b*_]} and *R*_*𝒥*_ = {[*x*_*j*_, *y*_*j*_]} are obtained by placing margins of size *l* (default *l* = 500 bp) around all breakpoints or novel junctions and merging overlapping intervals per allele.

Instead of actually sampling read pairs from the two alleles, we enumerate all possible combinations of starting position *p* and insert size *s* around breakpoints or novel junctions, respectively. In the following, we refer to these combinations as model read pairs. Using the variant description, we determine the model read pair’s mapping to the reference genome (i.e. its 5’-difference *d* and category *g*) and add it to the profile with weight *P*_Δ_(*s*). Formally, let the functions *d*^VAR^(*p, s*) and *g*^VAR^(*p, s*) perform the mapping and return *d* and *g*, respectively, for model read pairs from the variant allele. For model read pairs from the reference allele, the mapping is trivial. Here, *d* = *s* and we define the function *g*^REF^(*p, s*) that determines *g*. For the variant allele, we store model-read-pair counts for all possible combinations of *s, d*, and *g* in the matrix 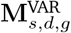. For the reference allele, we store read-pair counts of all possible combinations of *s* and *g* in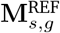 :

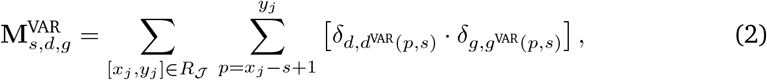

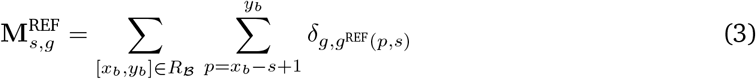

with *δ* denoting the Kronecker delta. Finally, we obtain allele-specific profiles by multiplying these matrices with the insert-size distribution:

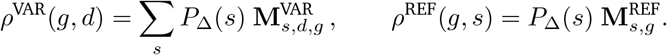

Further details on estimating insert size distributions and the mapping function can be found in the Supplementary Material Sec. 2.5 and 2.6.

#### 2.1.4 Genotype-specific read pair profiles

Analogous to the genotype being a combination of alleles a person carries at a given locus, the genotype profiles are determined by linear combinations of allele-specific profiles

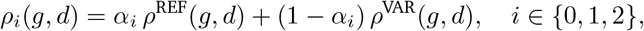

with coefficients *α*_0_ = 1 *−ϵ, α*_1_ = 1*/*2, and *α*_2_ = *ϵ*, where we introduce *ϵ ≪*1 for numerical stability. Note that the allele profiles *ρ*^REF^ and *ρ*^VAR^ are not normalized to unity in order to preserve the absolute number of read pairs expected from each allele.

Normalizing the above combinations *ρ*_*i*_, we obtain valid probability density functions 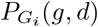 for our genotype profiles (Fig. 1D). These can be used to define the probability of observing a read pair *r* with 5’-difference *d*(*r*) and category *g*(*r*) given genotype *G*_*i*_:

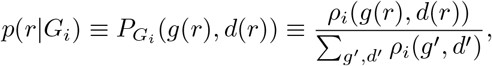

which completes our maximum-likelihood model of Eq. (1).

### 2.2 Genotype quality and certainty

The genotype quality is commonly defined as the ratio of the second largest to the largest likelihood (Nielsen et al., 2011). Most algorithms do not compute further measures of genotype confidence or the respective journal articles do not elaborate on them (Chiang et al., 2015; Chen et al., 2019; Niehus et al., 2021). However, sequencing is a stochastic process and the read pairs we observe will change between sequencing runs. This can lead to differences in genotype quality and ultimately the called genotype.

One tool that accounts for this randomness is BayesTyper, which models *k*-mer count distributions based on haplotype and diplotype frequencies and produces posterior genotype estimates via Gibbs sampling (Sibbesen et al., 2018). For the same purpose, we estimate a posterior distribution of genotype qualities (i.e. how much errors and randomness in the input data influence the genotype call) via bootstrap sampling of the set of read pairs ℛ (Fig. S11). We define the *genotype certainty* as the fraction of bootstrap samples in agreement with the original call and provide mathematical details in the Supplementary Material Sec. 2.8. Our genotype certainty condenses the posterior distribution into a robust estimate of how likely it is to obtain the same genotype upon repeated sequencing.

### 2.3 Genotype difficulty

Some variants are inherently more difficult to genotype than others due to their structure. For these, higher coverage is required to achieve reliable genotype calls. Our genotype profiles enable us to estimate the relative difficulty in genotyping different variants. The more the genotype profiles for a given variant differ from each other, the easier it will be to distinguish them based on observed read data, as a single read pair will on average provide more information about the underlying genotype. We quantify this using the Kullback-Leibler divergences (KLD) of two genotype profiles 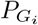 and 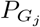 :

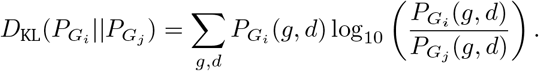

This can be interpreted as the expected gain in genotype quality upon observing a single read pair drawn from the distribution of the true genotype *G*_*i*_, when *j* is chosen such that 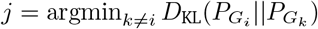.

We can use this to calculate the expected number *N*_*r*_(*v, Q*_*E*_, *i*) of read pairs required to genotype a variant *v* with genotype *G*_*i*_ with a certain quality *Q*_*E*_ and define the difficulty *D*(*v, Q*_*E*_) as

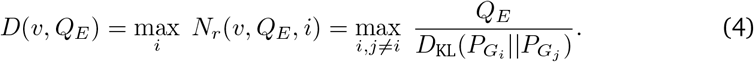

Our simulations below show that this quantity is a good indicator for the genotyping performance one can expect for a variant based on its structure. This may help in judging the reliability of genotype calls made on new data sets.

## 3 Results

We evaluated GGTyper on both simulated and real data. We compare GGTyper with Paragraph and BayesTyper, two graph-based tools that are able to genotype complex SVs when given VCF files containing reference and alternate allele sequences. Additionally, we tested Paragraph with variants supplied as graphs in a JSON file resulting in slightly different calls.

### 3.1 Simulations

To test our approach on simulated data, we created seven distinct data sets, each containing five diploid samples with 30x coverage. We used ART (Huang et al., 2011) for read pair simulation and BWA (Li and Durbin, 2009) for read alignment. Within each data set, we distributed 902 SVs of various types and sizes uniformly on chromosomes 20 and 21. We simulated canonical SV types (100 deletions, 100 inversions, 100 duplications, 100 tandem duplications, and 1 translocation) and six complex SV (csv) types (100 csvA, 100 csvB, 100 csvC, 100 csvD, 100 csvI, and 1 csvT). Detailed information on SV structures and sizes is available in the Supplementary Material Sec. 3.1. Samples within each data set share the same variants but have different haplotypes. In total, we evaluate 31 570 genotype calls.

Knowing the true genotypes for our simulated data, we compute the recall as the fraction of genotype calls passing all filters and the precision as the number of correct genotype calls divided by the number of genotype calls passing filters.

#### 3.1.1 Overall performance

Without any filtering of genotype results, GGTyper achieves a precision of 96.77 (*±* 0.16)%. Precision-recall curves indicate that genotype certainty is a better filtering metric than genotype quality (Fig. S1). Aiming for high precision, we set a minimum certainty of 0.9 and a minimum average read pair mapping quality of 40 as GGTyper’ default filters. As a result, we obtain a precision of 99.91(*±*0.02)% and a recall of 89.01(*±*0.30)%.

#### 3.1.2 Comparison to BayesTyper and Paragraph

Comparing the unfiltered precision of GGTyper to its competitors (Fig. 2), BayesTyper slightly outperforms GGTyper in terms of precision (98.37(*±*0.11)%) on simulated data. Paragraph’s performance is on par with the other methods for some variant types and far worse for others, leading to an overall precision of 78.70(*±* 0.21)% with graph input and 80.65(*±* 0.23)% with VCF sequence input.

**Figure 2.**
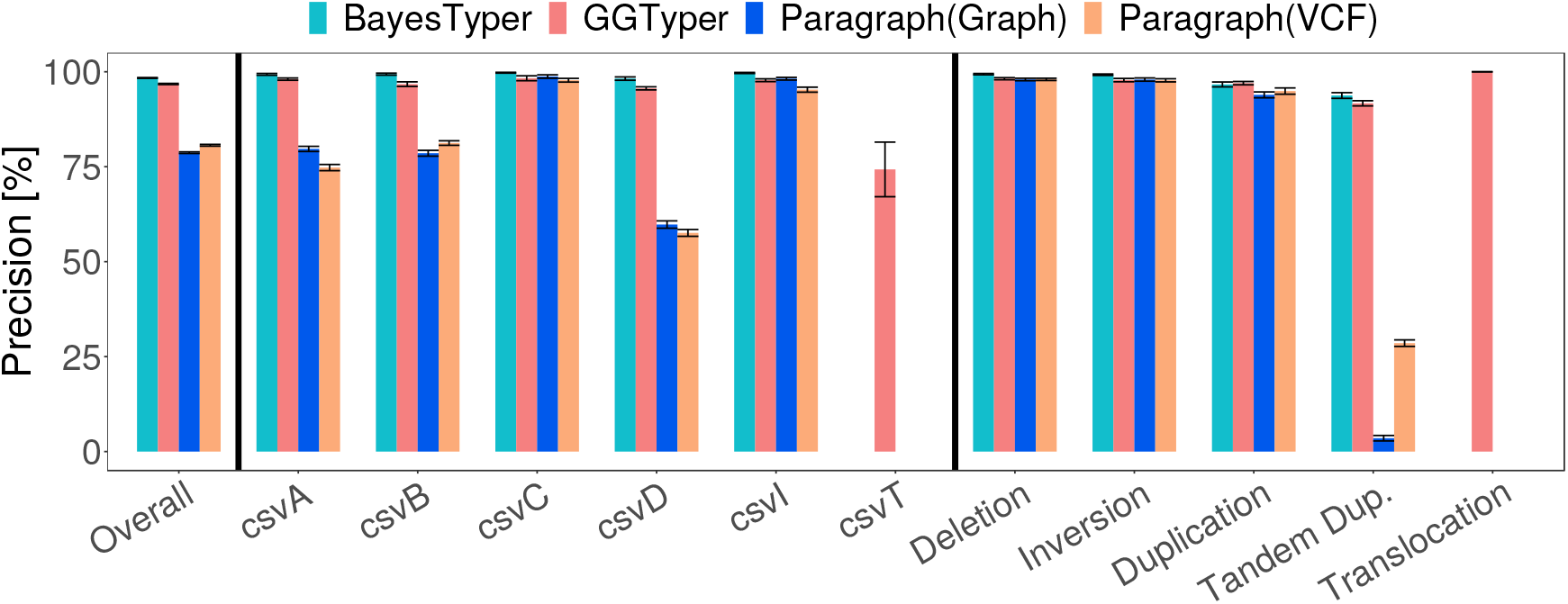
Precision of GGTyper, Paragraph and BayesTyper on simulated data by variant type using default filters of BayesTyper and no filters for GGTyper and Paragraph. Error bars indicate standard error across the seven simulated data sets.

Tandem duplications proved most difficult for all algorithms. Inter-chromosomal translocation variants (i.e. csvT and translocations) were excluded from genotyping with BayesTyper and Paragraph as they cannot be described in a single sequence. GGTyper’s difficulties in genotyping csvT can be entirely attributed to the input short read alignment since all incorrect calls are removed upon filtering by average mapping quality. Both GGTyper and BayesTyper show good performance when testing different coverages, with BayesTyper achieving a higher recall at almost identical precision (Fig. S2).

Both GGTyper and BayesTyper achieve higher precision and recall for complex than for canonical SVs (Fig. S2), indicating that complex SVs are in general not harder to genotype. This could be due to the fact that the majority of our simulated variants are located in regions with high mapping quality and their breakpoints are known exactly. Since complex variants are more likely to occur in repetitive regions (Sanchis-Juan et al., 2018), the high recall and precision shown here may not necessarily transfer to real data.

#### 3.1.3 Genotype difficulty predicts performance

We calculated the genotype difficulty (Eq. (4)) with *Q*_*E*_ = 100 for each simulated variant and examined the average difficulty per variant type. Fig. 3 indicates that both the genotype certainty and the precision decrease as the genotype difficulty of a variant type increases. Tandem duplications are most difficult on average, in accordance with the results described above. Furthermore, the figure demonstrates that the correlation between precision and genotype difficulty markedly decreases with increasing coverage, suggesting that our genotype difficulty reliably predicts how much data is needed to accurately genotype a given variant with a target certainty, with csvT being the only exception. The observed relationship between performance and difficulty could also be used to estimate whether a new data set has sufficient coverage to genotype a known variant with a target certainty.

**Figure 3.**
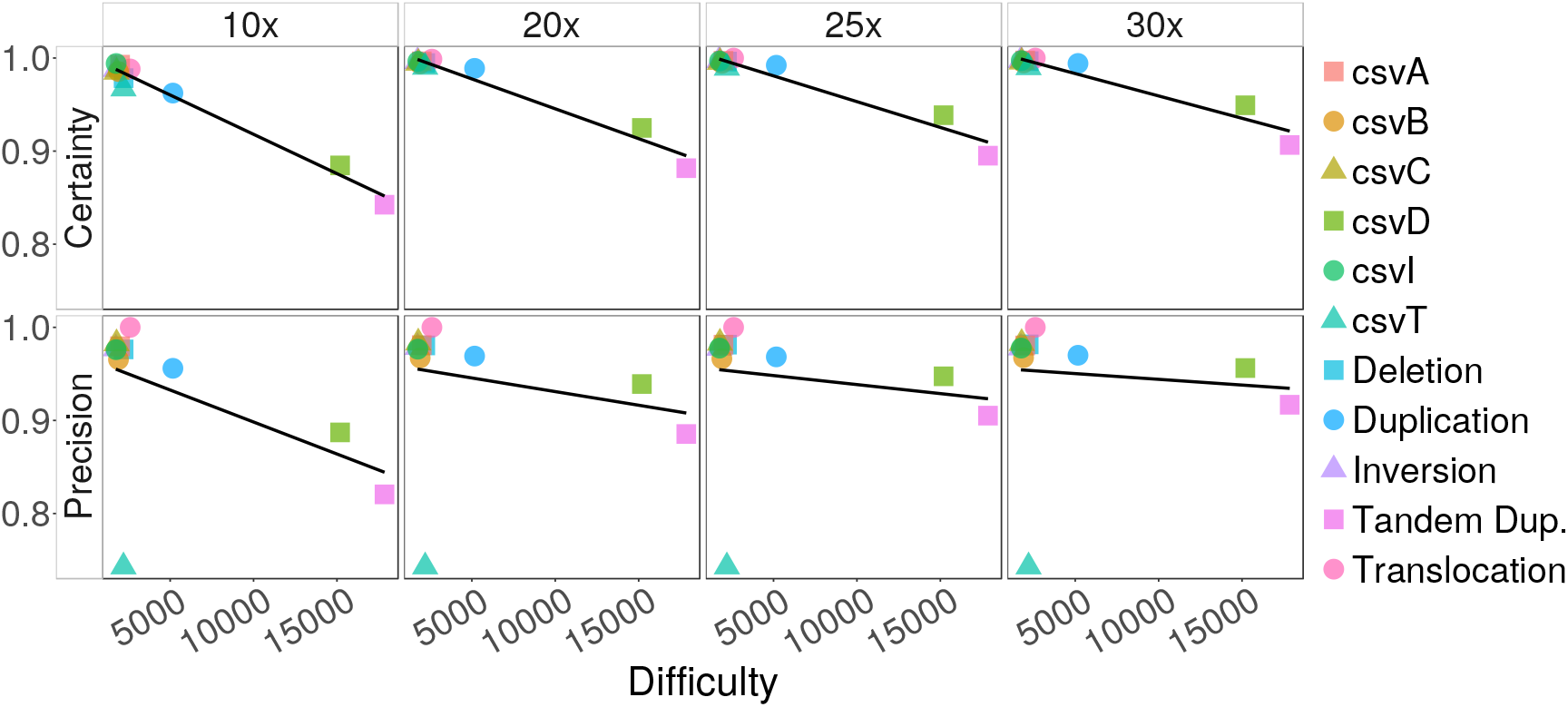
Genotype certainty and precision on simulated data depending on genotype difficulty (Eq. (4), Q_E_ = 100). Averages per variant type at 10x, 20x, 25x and 30x coverage are shown.

### 3.2 Real data

Our simulations show that GGTyper is able to genotype a wide variety of (complex) SVs with high accuracy. In order to evaluate how well the performance on simulated data translates to real-world applications, we genotyped a total of 20 manually curated complex SVs (V1, …, V20) from different sources (Supplementary Material Sec 3.2) in 7 829 Icelandic genomes (Gudbjartsson et al., 2015) comprising 3 340 trios and 199 genomes from the Polaris Diversity and Kids cohorts (https://github.com/Illumina/Polaris) comprising 49 trios from three continental groups.

Since benchmark sets for complex SVs are lacking, we rely on trio and population statistics to judge the accuracy of genotype calls. Following previous work (Niehus et al., 2021), we compute the Mendelian inheritance error rate (MIER), transmission rate (TR), and agreement with Hardy-Weinberg equilibrium (HWE) per variant. We calculate the MIER as the percentage of genotype calls in the children that violate the Mendelian rules of inheritance given the parents’ genotype calls. Focusing on trios where either the reference or the variant allele occurs only once in the two parents, we calculate the TR as the fraction of trios where the allele is passed on to the child. HWE is a model that connects allele frequencies to genotype frequencies within ideal populations. For each variant, we calculate expected genotype counts from the observed allele counts assuming HWE and compare expected to observed genotype counts using a *χ*^2^-test with *p* = 0.001 excluding children in trios.

#### 3.2.1 Icelandic data

We applied GGTyper to genotype our set of 20 complex SVs in all 7 829 Icelandic genomes. HWE, MIER and TR indicate high genotyping accuracy for most variants (Fig 4a). Of the 20 variants, 17 are in HWE. All 17 variants in HWE have a MIER below 1% and a TR very close to the expected value of 50%. Two of the three remaining variants show slightly elevated Mendelian inheritance errors (V20, V10) with minor elevations in the TR, while the third (V12) shows no inheritance errors at all but a TR close to 100%. This is due to an over-abundance of heterozygous variant calls caused by alignment difficulties in the variant region. In general, the MIER cannot detect this type of error and is known to be limited in measuring genotyping accuracy. However, the TR confirms incorrect genotyping of the variant. Without filtering, the overall TR across all variants is 51.12%.

**Figure 4.**
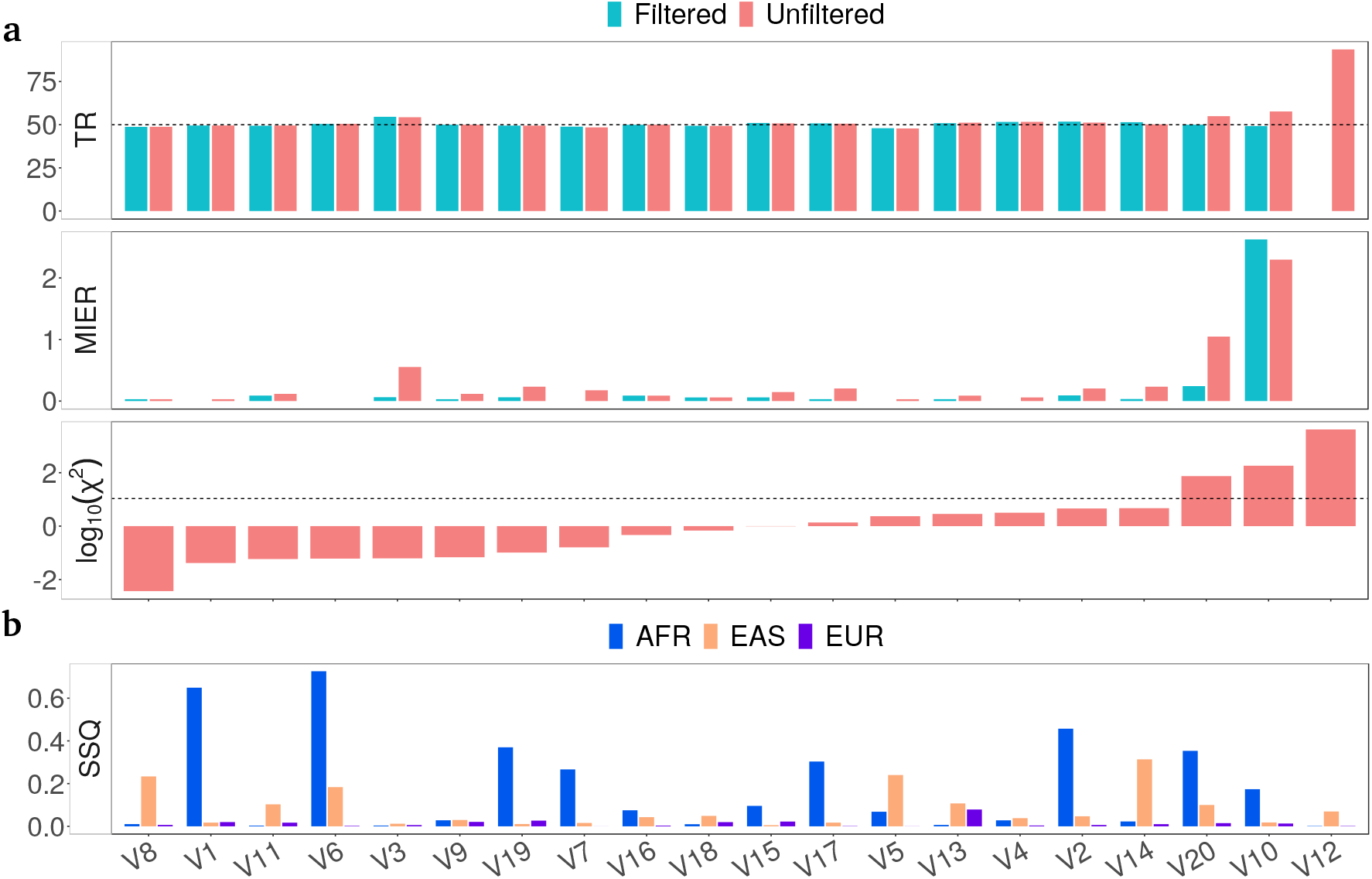
Population statistics on real data for 20 complex SVs. **a** TR and MIER in percent in 3 340 Icelandic trios and log-scaled χ^2^-value when testing for HWE in 4 289 Icelandic non-child genomes. Dashed lines mark the expected TR of 50% and the significance threshold (p = 0.001). We used default filters. **b** Sum-of-squares of the difference in genotype fractions between the Icelandic data and the three continental groups in the Polaris diversity cohort.

#### 3.2.2 Polaris data

We genotyped our set of 20 complex SVs in the 199 Polaris genomes using GGTyper, Paragraph and BayesTyper and measured computational performance on a subset (Table 1).

**Table 1.** Resource requirements for genotyping 20 complex SVs on Intel(R) Xeon(R) Gold 5115 CPU using 10 threads.

|  | GGTyper | Paragraph | BayesTyper |
| --- | --- | --- | --- |
| Wall time (10 genomes) | 2m 27s | 2m 48s | 5h 29m<sup>1</sup> |
| Memory (1 genome) | 0.66 GB | 3.1 GB | 29.4 GB |

Generating $k$ -mer bloom filters for BayesTyper took 13 hours.

In the 150 unrelated genomes, we calculated *χ*^2^-values determining which genotype predictions show HWE (Fig. 5). In the 49 trios, we calculated the TR. GGTyper’s variant calls are in HWE for again 17 variants but apart from V12 the variants failing HWE differ between data sets. Paragraph shows similar results to GGTyper, suggesting that the true genotype frequencies of V6 and V19 within the diversity cohort may not be in equilibrium. Contrary to what we observe in our simulations, BayesTyper performs worse than the other tools overall. Its default filters removed all non-reference genotype calls. Even after removal of all filters, no variant alleles were detected in any genome for 4 variants, no calls were available for V7, and another 5 variants fail HWE.

**Figure 5.**
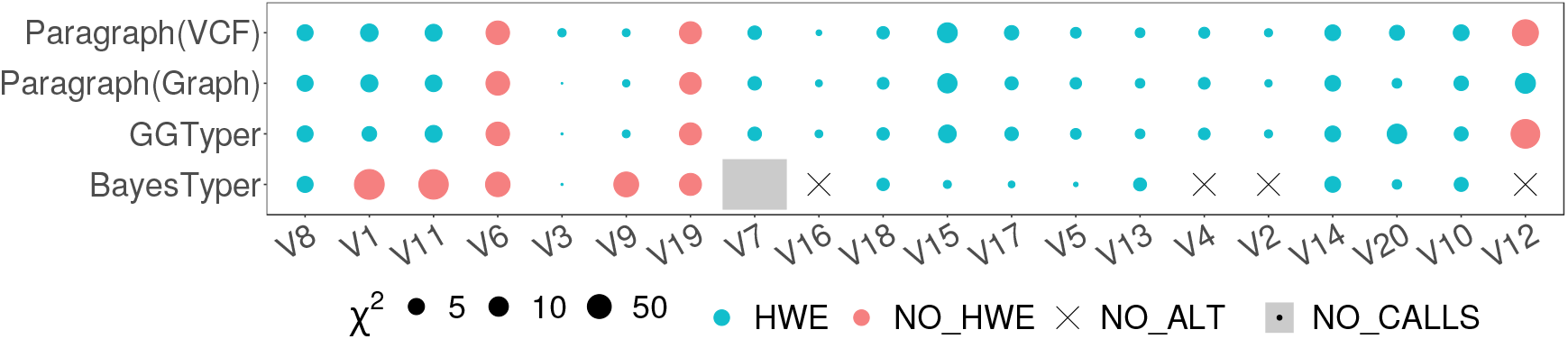
Status of HWE for 20 complex SVs in the Polaris diversity cohort. Point size indicates log-scaled χ^2^-value, colour indicates whether HWE could be rejected or not with p = 0.001. BayesTyper called 4 variants as homozygous reference in all samples (black crosses) and removed V7 when disabling all filters with BayesTyperTools filter.

GGTyper’s TR on unfiltered Polaris calls (51.05%) is very similar to that on the Icelandic data and to Paragraph (48.96% for Graph and 50.00% for VCF input). BayesTyper’s transmission rate (47.62%) is furthest from the theoretical value. We note that the variant without call and the 4 variants, where BayesTyper does not call variant alleles, do not contribute to the transmission rate. Fig. 6 shows the TR for all three tools with different degrees of filtering. When increasing the filter strength, BayesTyper’s transmission rate fluctuates around 50% while GGTyper and Paragraph diverge from 50% in opposite directions. Overall, GGTyper’s results stay closer to the expected value for filtering down to about 50% of the trios. Filtering GGTyper’s calls by genotype certainty leads to less biased results in terms of TR but the percentage of trios that can be filtered is limited as *𝒞 ≤* 1 by definition. GGTyper’s performance is very similar on the two different data sets, with the Icelandic results being more stable due to the larger number of trios available. Overall, we can conclude that GGTyper consistently performs on a high level across data sets.

**Figure 6.**
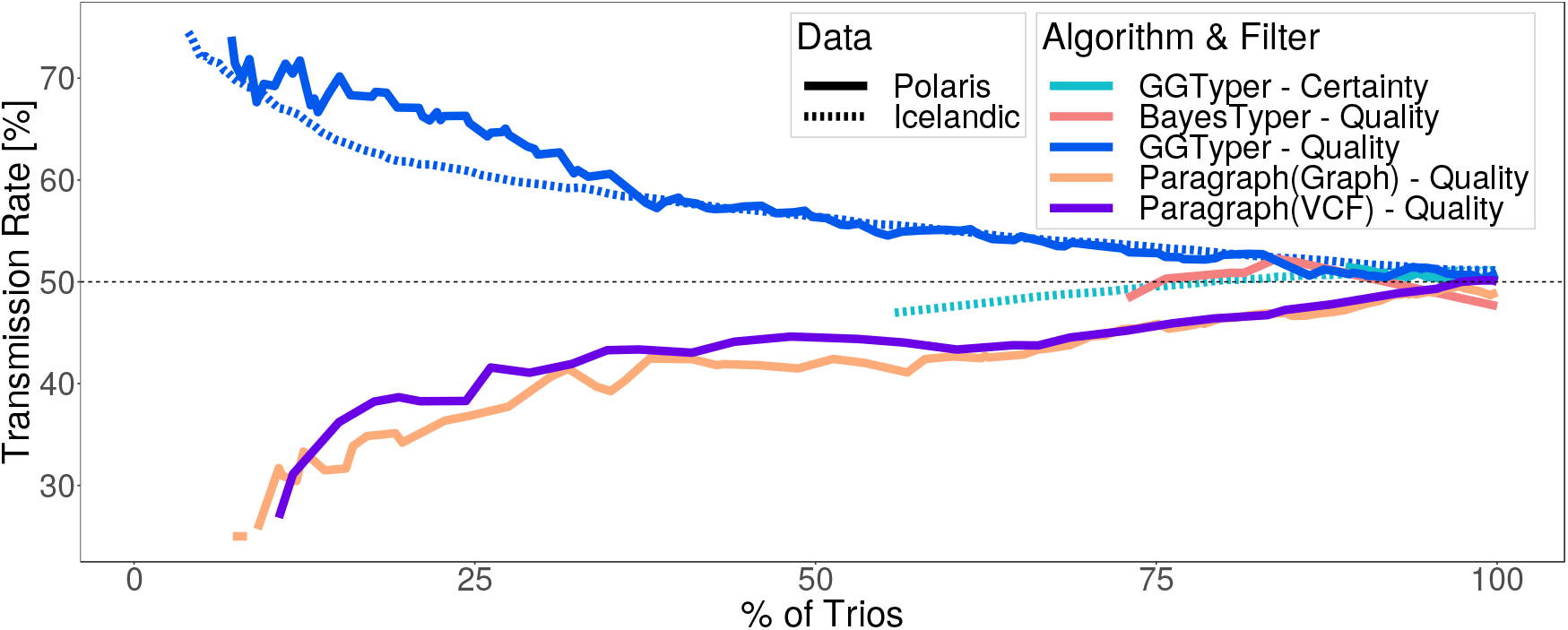
TR across 20 complex SVs in both Icelandic and Polaris data for GGTyper, Paragraph with graph or VCF input, and BayesTyper, filtered by decreasing genotype quality or certainty. Only 15 variants contribute to BayesTyper’s TR.

Testing for HWE shows clear differences between the Icelandic and Polaris data sets (Fig. S6). This may be due to the much smaller size of the Polaris diversity cohort, but can also be attributed to the heterogeneous composition of that data set. Calculating the difference in genotype frequencies between each of the three continental groups in the Polaris data and the Icelandic genomes reveals significant differences (Fig. 4b). As expected, the Icelandic genomes are most similar to the European genomes.

## 4 Discussion

GGTyper implements a new algorithm that serves the available masses of short-read sequencing data by efficiently and accurately genotyping known complex SVs. Our approach goes beyond many previous genotyping approaches that use a binary read pair classification into reference-supporting and variant-supporting. It circumvents the problem of combining possibly conflicting genotype calls of several novel junctions into a single genotype call. Our normalization of genotype profiles takes care of unequal numbers of breakpoints and novel junctions between alleles. In addition, our genotype profiles enable definition of a new measure that judges an SV’s genotyping difficulty before analysing data.

While the detection of complex SVs in short-read data is challenging, our simulation confirms that complex SVs are not inherently more difficult to genotype than canonical SV types. On simulated data, GGTyper achieves high precision and recall when genotyping SVs with a wide range of structures. We suspect that Paragraph’s low precision and its inability to genotype tandem duplications is due to the fact that it uses only reads with unique graph alignments. We attribute BayesTyper’s almost perfect performance in the simulation to the alignment-free approach. In contrast to BayesTyper, GGTyper and Paragraph take read alignments as input and alignment quality has a significant impact on genotyping performance. GGTyper achieves high precision when filtering for variants with high mapping quality of corresponding read pairs, at the cost of a notable drop in recall. Using a more complete reference like T2T-CHM13 (Nurk et al., 2022) was already shown to readily improve genotyping (Aganezov et al., 2022). For our assessment, a realignment of all previously aligned data was computationally unaffordable.

As opposed to GGTyper’s consistently high performance, BayesTyper’s accuracy on simulated data does not translate to real data. A potential explanation for this is imprecise specification of breakpoint sequences (e.g. due to micro-insertions or microhomologies), which cause the *k*-mer based approach to fall behind. Our approach is by design robust against small inaccuracies in the variant specification. Compared to the simulation results, Paragraph performs far better on real data and is mostly on par with GGTyper in terms of transmission rate. Paragraph’s varying performance for different types of variant structures in our simulations suggests that our set of 20 complex SVs on real data may not cover a wide enough range of variant structures. Application to larger sets of variants (e.g. obtained by comparing telomere-to-telomere assemblies) is left for future work.

Our approach generalizes to multi-allelic SVs (i.e. loci with more than one alternate allele including low-copy tandem repeats) and the implementation has already been extended but not yet benchmarked. In addition, GGTyper is currently unable to recognize alleles that are not provided in the input variant description. This may lead to incorrect genotype calls and needs to be addressed in the future. Another limitation of our approach is its inability to genotype variants containing non-reference sequence insertions. Although many complex variants are simply rearrangements of existing genomic segments (Collins et al., 2017), we previously observed non-reference sequence involved in complex SVs (Kehr et al., 2017) and strive to extend GGTyper accordingly.

Despite these limitations, GGTyper fills an important gap when it comes to reliably genotyping known, breakpoint-resolved SVs until long-read data has become as readily available as short-read data and pangenomes supersede linear reference genomes.

## 6 Author contributions statement

T. M., R.L. and B.K. conceived the method. T.M. implemented GGTyper and conducted the experiments on simulated and Polaris data. B.V.H. conducted the experiments on Icelandic data. T. M., R.L. and B.K. analysed the results and wrote the manuscript. All authors reviewed the manuscript.

