## Supplementary Material for "GGTyper: genotyping complex structural variants using short-read sequencing data"

March 14, 2024

### 1 Supplementary Figures

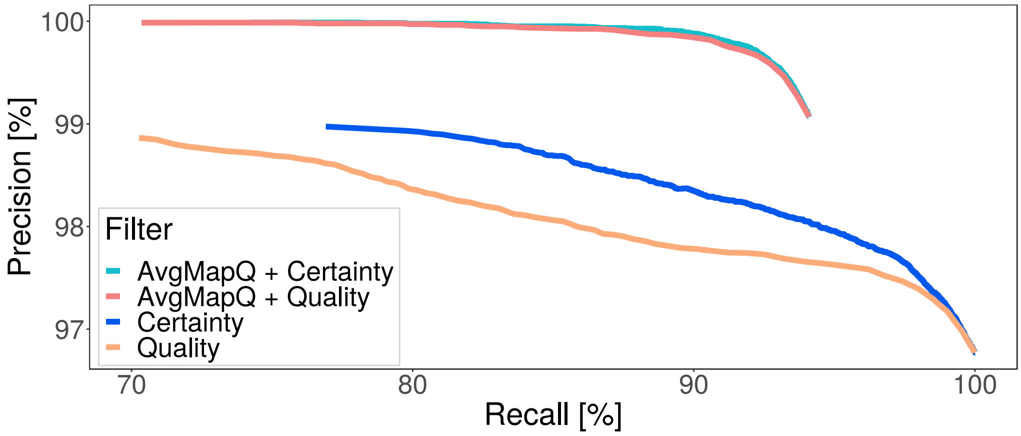

Figure S1: Precision-recall curve for GGTyper on simulated data obtained by filtering for increasing genotype certainties or genotype qualities. The minimum average mapping quality (AvgMapQ) was fixed to 40 when applied as filter.

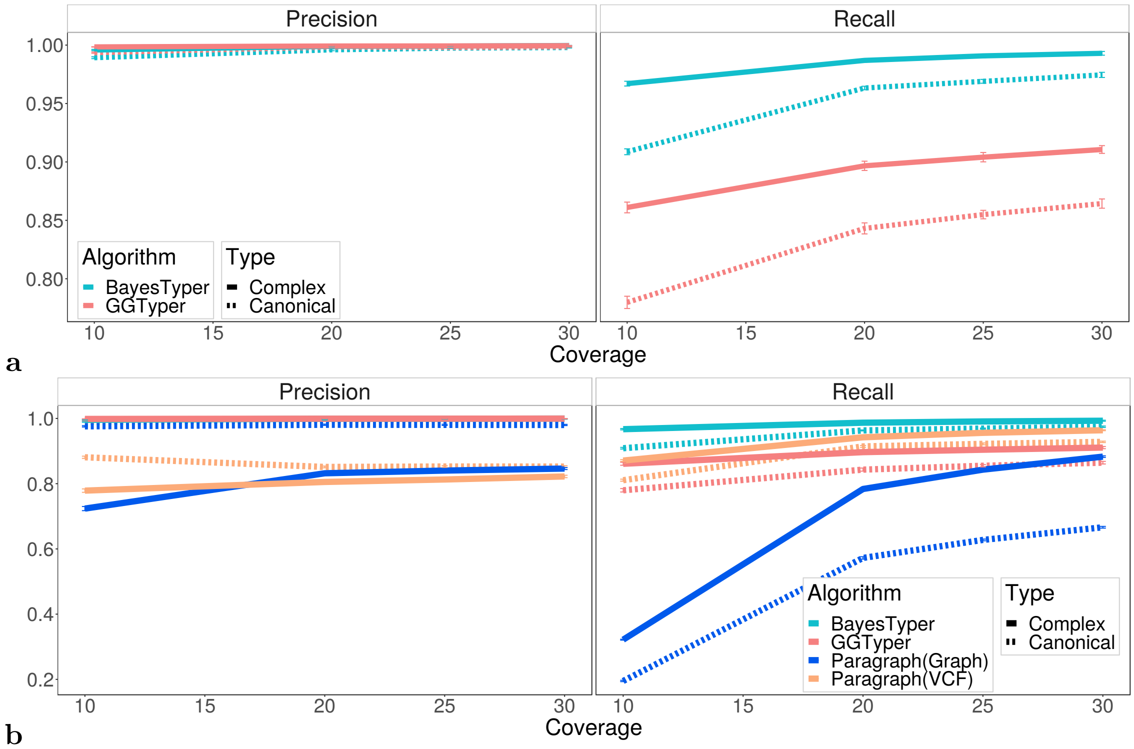

Figure S2: **a)** Precision and recall of GGTyper and BayesTyper for data simulated at 10x, 20x, 25x, and 30x coverage split by complex and canonical SVs. GGTyper's genotype calls were filtered by average mapping quality ($\geq 40$) and genotype certainty ($\geq 0.9$). BayesTyper genotype calls were filtered by genotype quality ($\geq 20$) and otherwise default filters. **b)** Same plot including Paragraph.

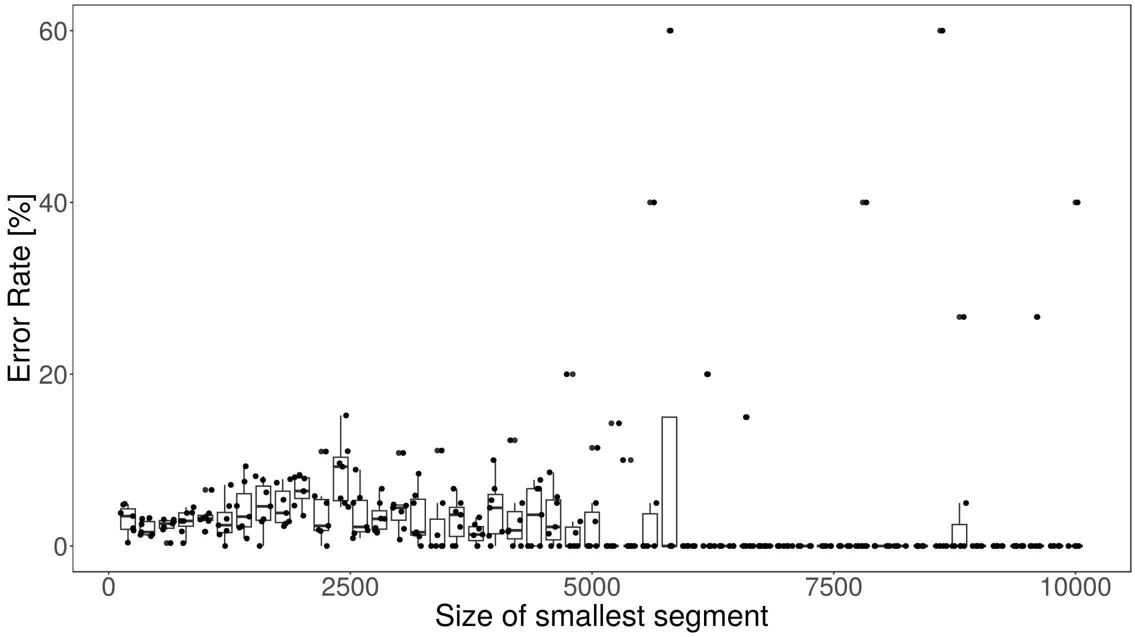

Figure S3: Error rate of GGTyper's genotype calls on simulated data depending on the size of the smallest DNA segment involved in the genotyped SV. Error rate is plotted for every data set in each size bin. One bin has size 200 bp.

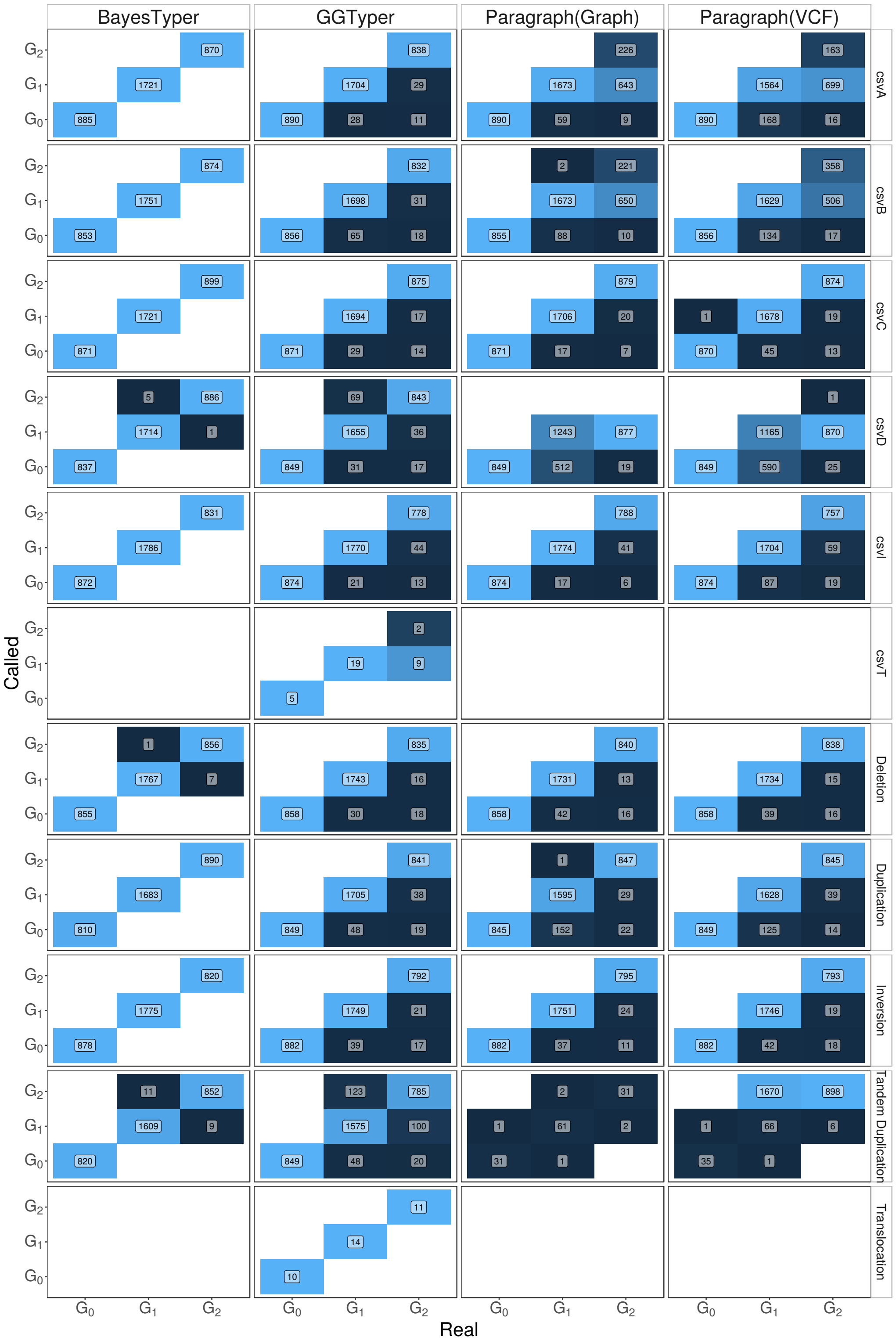

Figure S4: Number of genotype calls $N$ (unfiltered) for each combination of simulated ground-truth and called genotype at 30x coverage. Colour (dark blue: 0, light blue: 1) indicates fraction of $N$ relative to the total number of calls for the respective ground-truth genotype for the given variant type.

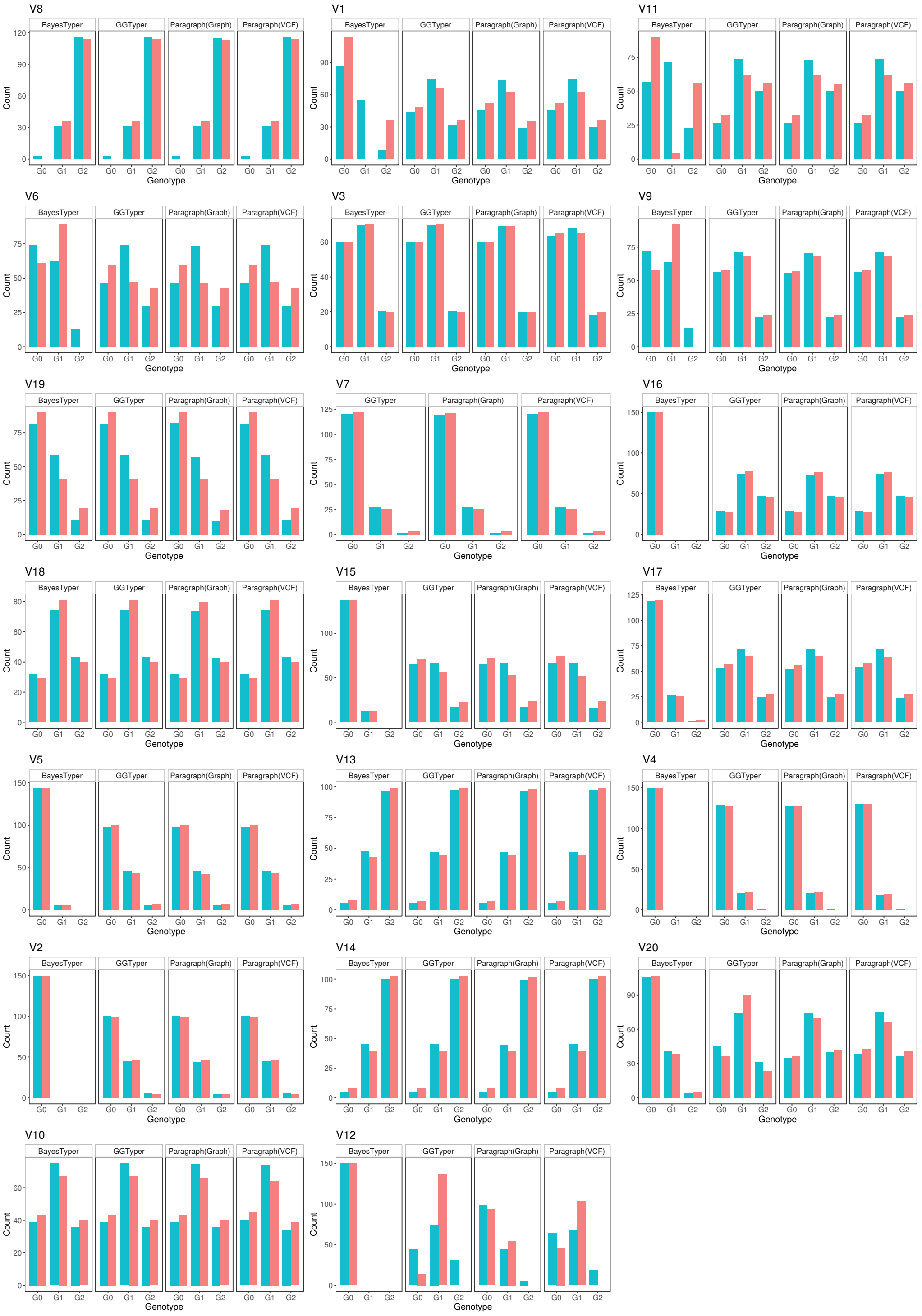

Figure S5: Observed and expected genotype counts for unrelated samples from Polaris Diversity cohort as obtained by GGTyper, Paragraph (VCF or Graph input) and BayesTyper.

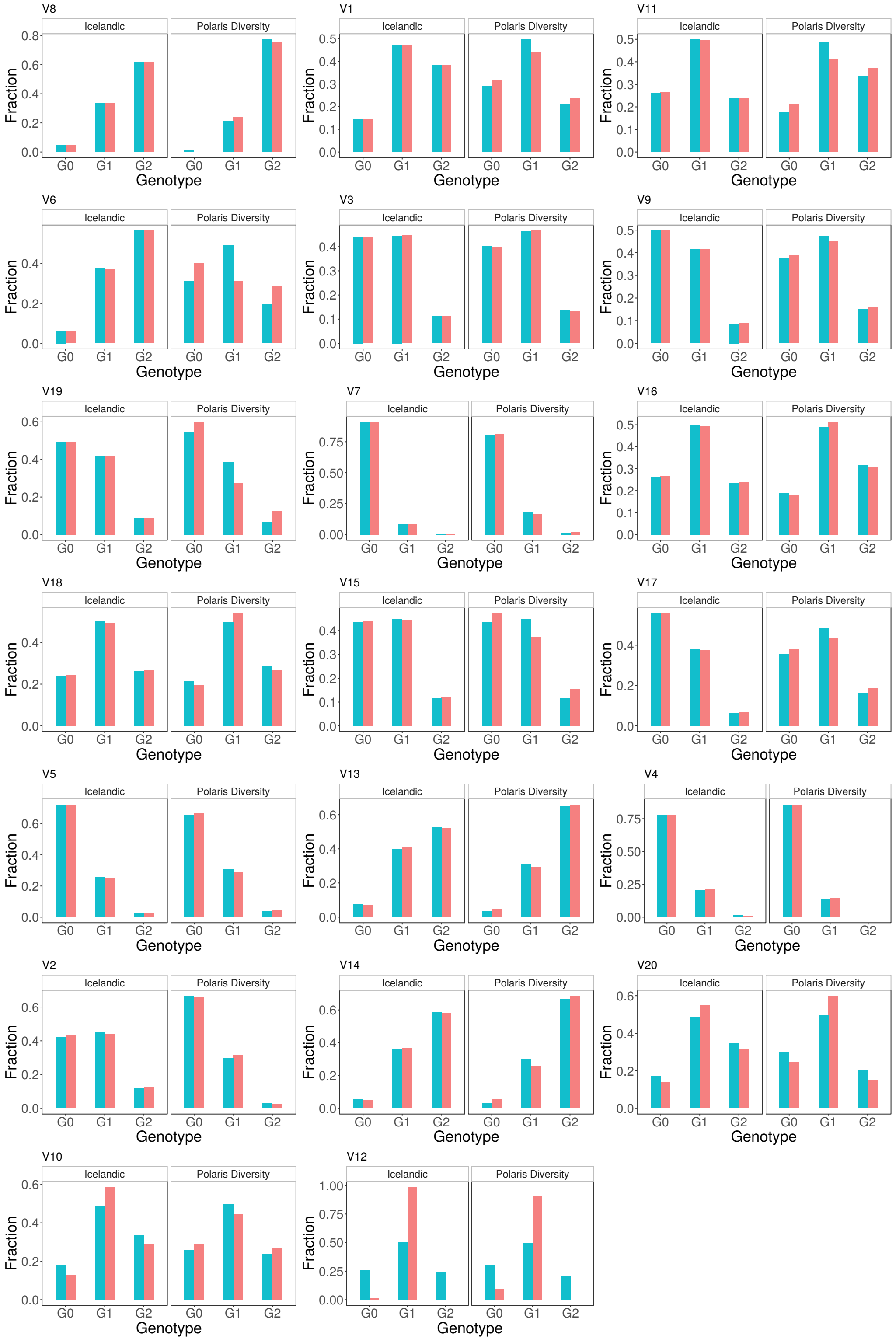

Figure S6: Observed and expected genotype counts for unrelated samples from Polaris Diversity cohort and unrelated Icelandic samples as obtained by GGTyper.

### 2 Supplementary Methods

#### 2.1 Variant description

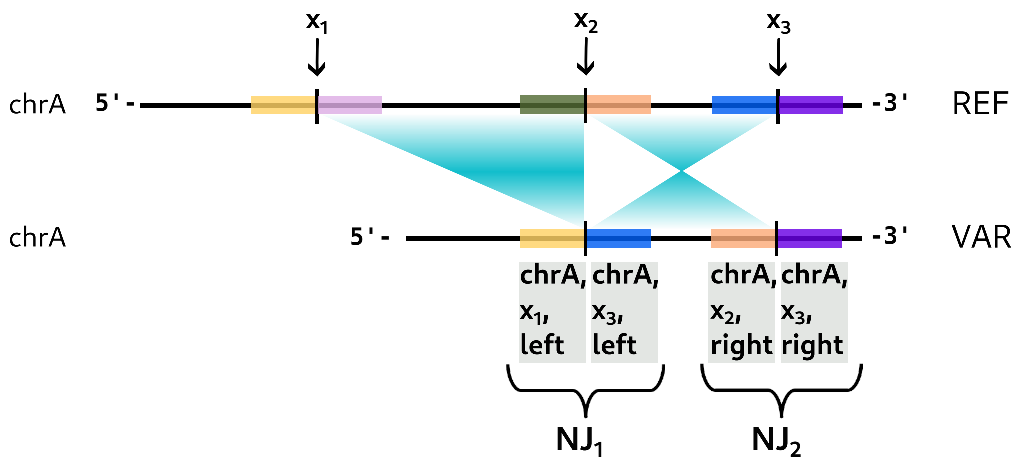

Figure S7: Representation of a complex variant containing a deletion and an inversion. The positions of the reference breakpoints on chromosome A are denoted by $x_1$, $x_2$, and $x_3$. $\mathrm{NJ}_1$ and $\mathrm{NJ}_2$ are the two novel junctions that describe the variant sequence. The information contained in the junctions is shown in the grey boxes. Coloured squares are used to visualize areas to the left and right of the breakpoints and where they end up on the variant.

Complex structural variants occur when DNA breaks at multiple positions and the resulting fragments are joined in a new configuration that differs from the original sequence. Disregarding the possibility of micro-insertions or other side effects of these mechanisms, this knowledge can be used to create a framework for the description of arbitrary variants. The positions where the reference sequence breaks are referred to as *reference breakpoints*. Simply put, the DNA repair mechanisms of the cell re-join the ends of fragments resulting from such breaks in order to repair the DNA strands. If the fragments are not joined in the original configuration, this creates so-called *novel junctions*. Each novel junction can be described by the ends of the two fragments that are newly joined. Considering a single novel junction, these two fragments may either be in their original orientation or in reverse orientation.

The 5'-fragment of the novel junction is in reverse orientation if the joined end originates in 3'-direction of a reference breakpoint, the 3'-fragment of the junction must have been reversed if the joined end originates in 5'-direction of a reference breakpoint. Consequently, both the 5'- and 3'-segment of the novel junction are described by a reference breakpoint and the direction (5'/left or 3'/right) of the joined sequence relative to the breakpoint. Figure S7 visualizes this for a complex SV. Note that a similar way of description using so-called *breakends* is part of the VCF format in order to describe arbitrary variants [1].

The sequence of a variant allele can be directly recreated from the supplied ordered set of novel junctions $\mathcal{J}$ and the set of reference breakpoints $\mathcal{B}$ can be obtained simply by splitting all novel junctions and removing potential duplicated breakpoints.

##### Description of variant in Fig. S7 in JSON-format

```
"cxSV1" : {                         # variant name
  "VAR" : {                         # name of the first alternate allele
    "chrA" : {                      # name of the affected chromosome,
                                    # must match bam file header
      "1" : {                       # first junction on altered chromosome
        "rNameLeft": "chrA",        # Name of chromosome of origin
                                    # of segment on 5'-side of junction
        "rNameRight": "chrA",       # Name of chromosome of origin
                                    # of segment on 3'-side of junction
        "xLeft": x1,                # First position on chromosome of
                                    # origin of 5'-segment
        "xRight": x3,               # First position on chromosme of
                                    # origin of 3'-segment
        "directionLeft": "left",    # 5'-segment originates to the left (5')
                                    # of pos. "xLeft" on chr "rNameLeft"
        "directionRight": "left"    # 3'-segment originates to the left (5')
                                    # of pos. "xRight" on chr "rNameRight"
      },
      "2" : {                       # second junction
        "rNameLeft": "chrA",
        "rNameRight": "chrA",
        "xLeft": x2 + 1,
        "xRight": x3 + 1,
        "directionLeft": "right",
        "directionRight": "right"
      }
    }
  }
}
```

#### 2.2 5'-difference

With $x_{\ell,5'}$ and $x_{r,5'}$ denoting the $5'$-ends of the left (closer to $5'$-end of reference chromosome) and right read in a pair, respectively, we define

$$d = z_1 - z_0 + 1, \qquad (z_1, z_0) = \begin{cases} (x_{\ell,5'}, x_{r,5'}) \text{ if orientation } RF, \\ (x_{r,5'}, x_{\ell,5'}) \text{ else.} \end{cases} \tag{1}$$

Using this definition, the 5'-difference of a read pair becomes negative for orientation $RF$ and positive for all other orientations (see Fig. S8).

#### 2.3 Detection of read-pair properties

##### 2.3.1 Split read pairs

A read pair is marked as split if any of the reads in the pair is split at a novel junction of the variant allele. Ideally, such a read would result in two read fragments mapped in the vicinity of separate reference breakpoints. However, the observed alignment data often

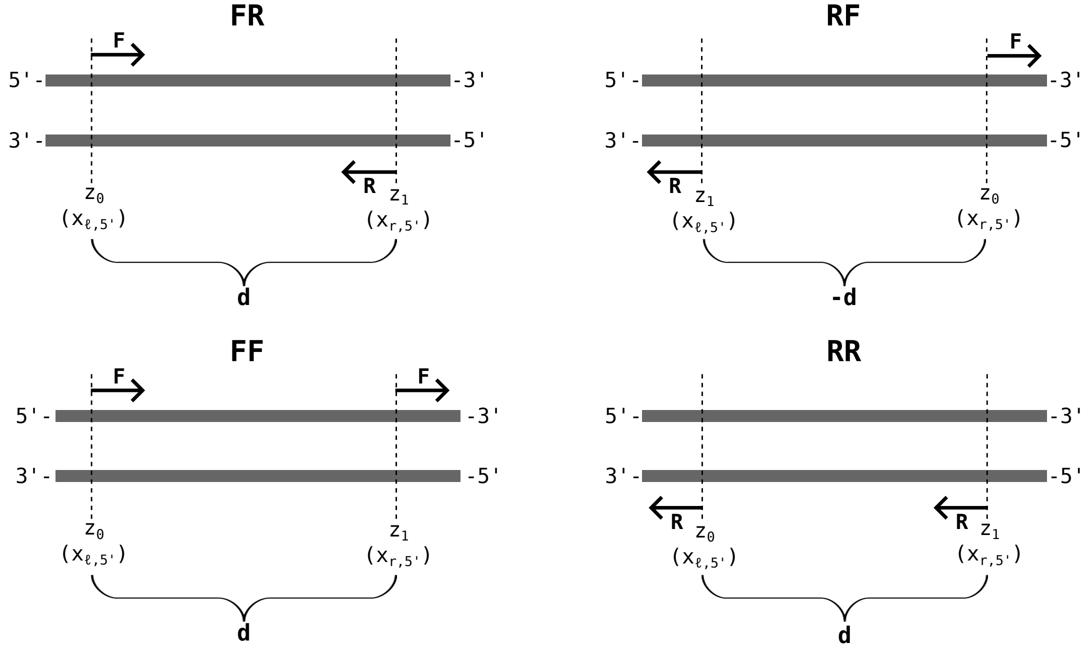

Figure S8: Definition of 5'-difference $d$ depending on pair orientation.

contains only one fragment of the split read. In order to avoid missing such reads, we rely solely on the primary alignment of each individual read and mark it as a split read if

1. it is clipped by at least 20 bp at a reference breakpoint,
2. aligns at a maximum distance of 20 bp from the breakpoint,
3. and does not overlap the breakpoint by more than 5 bp.

Alternatively, split reads may also be detected from gapped alignment of a record, which can occur for small deletions (not much larger than 50 bp when using BWA-mem with default settings). For each read pair, we determine the combination of novel junctions at which the individual reads are split. The maximum size of that set according to our definition of split reads is 4, since we consider only the primary alignment for each read and detect splits via clipping of its ends, which means that one read can be split at no more than two novel junctions.

##### 2.3.2 Spanning read pairs

Spanning read pairs are read pairs containing at least one read that spans a breakpoint on the reference allele. Their detection is straightforward. In order to stay robust with respect to small alignment errors and consistent with the definition of split reads, a read is only classified as spanning a breakpoint, if at least 20 bp of the read align on both sides of the breakpoint. For each read pair, we determine the combination of reference breakpoints that are spanned by the individual reads.

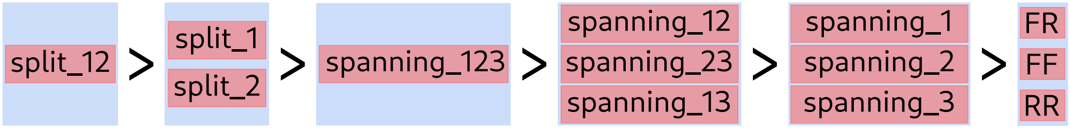

Figure S9: Ordering of created categories for a complex variant consisting of a deletion and an inversion (3 breakpoints, 2 novel junctions). Categories of the same ranking (in the same blue box) are mutually exclusive. A read pair is assigned to the highest ranking category corresponding to a subset of observed properties.

##### 2.3.3 Orientation

Orientation is simply defined as the combination of the individual orientations of the two reads in a pair. Due to the ability of the 5'-difference to distinguish between $FR$ and $RF$ orientation (see Sec. 2.2), we only assign read pairs to one of $\{FR, FF, RR\}$.

#### 2.4 Categories based on properties

In order to avoid a large number of potentially very small categories, we let some classes of properties take precedence over others. First of all, if a read pair is already marked as containing split reads, we ignore spanning or orientation information. On the other hand, if a read pair is not marked as split but instead marked as containing breakpoint-spanning reads, we do not take orientation information into account. Finally, if a read pair contains neither split nor breakpoint-spanning reads, we categorize it by its orientation.

Only the properties *FR, FF, RF* are mutually exclusive. For split and spanning read pairs, a read pair is always categorized by its entire set of relevant junctions or breakpoints. For example, consider a read pair that contains reads both split at novel junction 1 and novel junction 2. This read pair is then assigned to neither category *split_1* nor *split_2* but instead to category *split_12* (see Fig. S9).

#### 2.5 Library-specific insert size distribution

We assume that fragments created from the variant and the reference allele follow the same library-specific distribution of insert sizes. For read pairs that are in accordance with the reference genome, the 5'-distance is equivalent to the insert size of the fragment. We therefore determine the empirical distribution of insert sizes $P_\Delta$ by sampling 80 non-overlapping regions of length 25000 bp, filtering read pairs for correct alignment, specifically

- $0 <$ insert size $< 1000$,
- both reads mapped as primary records,
- both reads passing the aligner's QC,
- both reads have mapping quality $\geq 40$,
- no clipping or deletions,

and storing the insert sizes in a histogram. After smoothing with a Gaussian kernel, the histogram is normalized to 1 to obtain $P_\Delta$. Due to the random selection of genomic regions, sample profiles and final genotype likelihoods differ slightly across repeated runs.

However, the number and size of regions is large enough that these fluctuations do not negatively affect reproducibility.

If the BAM file contains only a small section of reads, GGTyper can use all aligned read pairs in the file in order to create the insert size distribution.

#### 2.6 Mapping

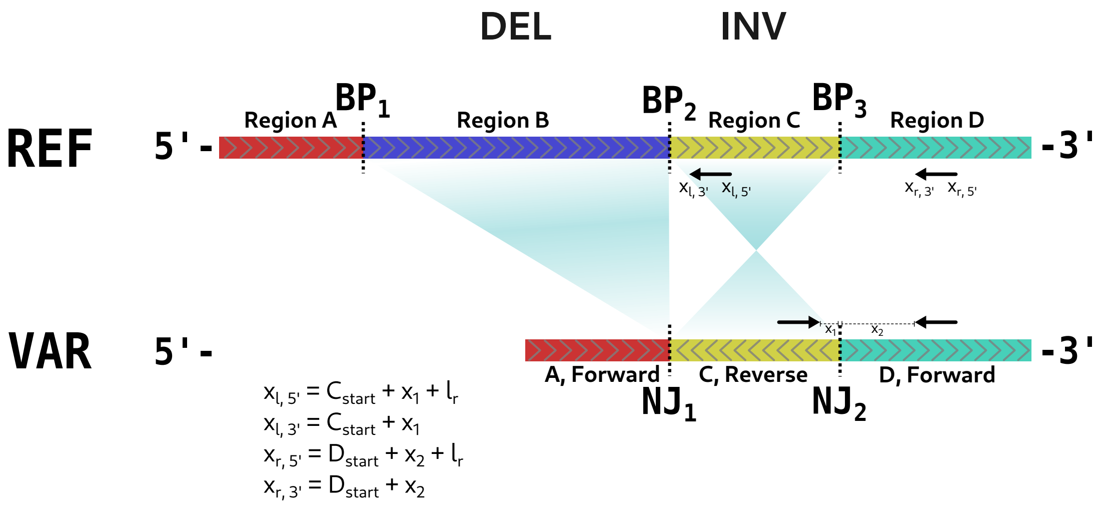

Figure S10: Variant allele as re-arrangement of reference DNA segments. $x_{\ell,5'}, x_{\ell,3'}, x_{r,5'}$ and $x_{r,3'}$ are coordinates of the read pair with respect to the reference (after mapping). Regions are specified by *chromosome:start-end*, where *start* and *end* are coordinates on the reference chromosome and *start* $<$ *end*. This figure shows the calculation of read-pair coordinates in a simple scenario without clipping. With clipping, either $l_r$ (read length) or the positions relative to the regions $(x_1, x_2)$ must be adjusted accordingly. After mapping a read is in forward orientation $(F)$ if its $5'$-coordinate is smaller than its $3'$-coordinate and reverse $(R)$ otherwise.

Our functions $d^{\mathrm{VAR}}(p, s)$ and $g^{\mathrm{VAR}}(p, s)$ contain a mapping from the variant to the reference allele. Since the variant allele is just a re-arrangement of the reference allele, mapping of simulated read pairs to the reference is straightforward. We only need to determine the 5'- and 3'-coordinates of the reads in a pair relative to the reference regions that make up the variant allele while considering the orientation of the regions and the orientations of the two reads on the variant allele. Reads that overlap a novel junction are clipped at these junctions, retaining only the largest fragment, before their reference coordinates and orientation after mapping are determined as illustrated in Fig. S10.

#### 2.7 Calculation of histograms

While the actual calculation of the allele-specific read-pair profiles is slightly more complex due to the creation of the matrices $\mathbf{M}^{\mathrm{VAR}}$ (Eq. (2) in the main text) and $\mathbf{M}^{\mathrm{REF}}$ (Eq. (3)

in the main text), the pseudocode for a naive calculation of the profiles is very simple, see Alg. 1.

**Algorithm 1** Read-pair profile calculation

```
for every allele a do
    initAlleleHistogram()
    for every insert size s do
        startPositions = getOverlapPositions(a, s)
        for x in startPositions do
            p = P_Δ(s)
            mappedPos = getReferenceCoordinates(x, s)
            attributes = getMappedAttributes(mappedPos)
            addToAlleleHistogram(attributes, p)
        end for
    end for
end for
```

#### 2.8 Bootstrapping of observed read pairs

With the log-likelihood ratio $\mathcal{Q}_{i,j}(\mathcal{R}) = \log_{10}(\mathcal{L}_i(\mathcal{R})/\mathcal{L}_j(\mathcal{R}))$ of two genotypes $G_i$ and $G_j$, the *genotype quality* is defined as

$$\mathcal{Q}(\mathcal{R}) = \min_{l \neq k} \mathcal{Q}_{k,l}(\mathcal{R}), \qquad k = \text{argmax}_{i \in \{0,1,2\}} \mathcal{L}_i(\mathcal{R}). \tag{2}$$

However, in order to accurately judge the reliability of a genotype call, the posterior distribution of $\mathcal{Q}$ has to be considered.

To this end, we determine the best and second best genotypes $k$ and $l$ based on $\mathcal{R}$. We then create a large number of bootstrap samples from the set of read pairs $\mathcal{R}$, and calculate $\mathcal{Q}_{k,l}(\mathcal{S})$ (for fixed $k$ and $l$) within each individual bootstrap sample $\mathcal{S}$. The distribution of these log-transformed likelihood ratios then provides an estimate for the posterior distribution of $\mathcal{Q}(\mathcal{R})$, which can be used to determine confidence intervals (see Fig. S11 for an example).

We furthermore define the *genotype certainty* as the fraction of bootstrap samples in agreement with the genotype call based on the original set of reads $\mathcal{R}$, that is

$$\mathcal{C}(\mathcal{R}) = \frac{1}{|\mathscr{S}|} \sum_{\mathcal{S} \in \mathscr{S}} \Theta \left( \min_{i \neq k} \mathcal{Q}_{k,i}(\mathcal{S}) \right), \tag{3}$$

where $\Theta$ denotes the Heaviside step function and $\mathscr{S}$ is the set of bootstrap samples. Note that in Eq. (3) the index $k$ is fixed based on $\mathcal{R}$, see Eq. (2).

#### 2.9 Implementation

Our algorithm is implemented in the program GGTyper (`https://github.com/kehrlab/Complex-SV-Genotyping`). It is written in C++, relying on *Seqan* [2] and *htslib* [3] for handling of sequence and alignment data and *Eigen* for fast matrix operations. Details on input files and parameters as well as the output format may be found on our GitHub page. The runtime for GGTyper scales linearly with the number of samples and the number of variants. Memory requirements are generally low with < 100 MB per thread.

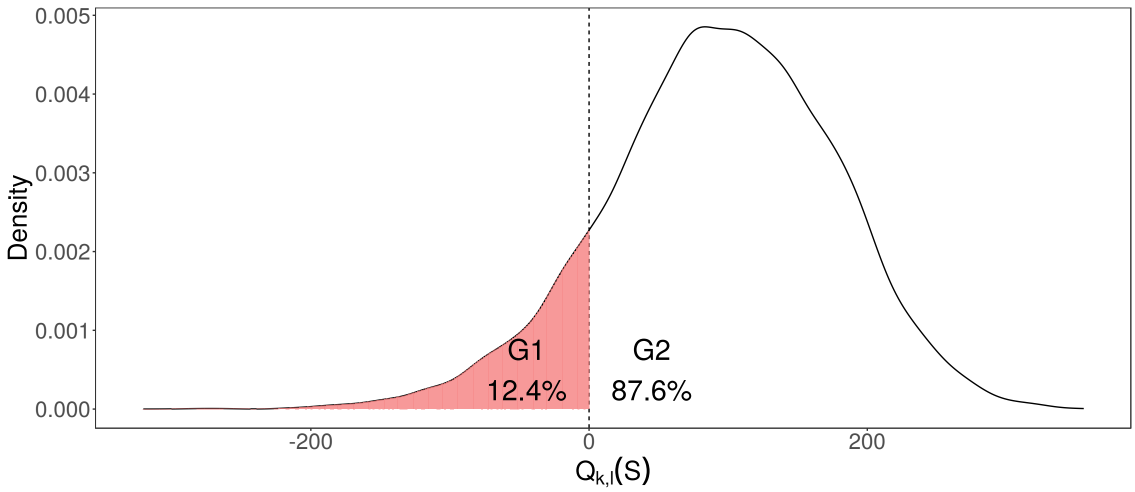

Figure S11: Example of a posterior distribution of the quality $\mathcal{Q}(\mathcal{R})$, see Eq. (2), obtained by bootstrapping of the set of observed read pairs $\mathcal{R}$ and resulting in $\mathcal{C} = 0.876$.

##### 2.9.1 Matrix formulation

Using the matrices $\mathbf{M}^{\mathrm{VAR}}$ (Eq. (2) in the main text) and $\mathbf{M}^{\mathrm{REF}}$ (Eq. (3) in the main text), variant profiles may be calculated and stored independently of the library distributions of the individual samples. This means we need to simulate possible read pairs and their attributes only once for each variant and reduces the actual creation of genotype profiles to matrix operations, which are efficiently implemented using *Eigen's* sparse matrices.

##### 2.9.2 Structure

GGTyper is split into three distinct parts.
*Profile-samples* calculates library distributions (see Sec. 2.5) and stores them together with read length and metadata as sample profiles in binary files.
*Profile-variants* counts, for all alleles of the given variants and for every possible insert site in a given set of library distributions, the number of possible read pairs created with specific attributes. These numbers are stored in allele-specific matrices $\mathbf{M}^{\mathrm{VAR}}$ and $\mathbf{M}^{\mathrm{REF}}$ and written to binary files as variant profiles (together with corresponding variant metadata).
*Genotype* calculates genotype profiles for every combination of variant profile and sample profile and determines the most likely genotype from the BAM files belonging to the sample profiles.

#### 2.10 Comparison tools

##### 2.10.1 Paragraph

We use the static build of Paragraph [4] v2.4a (https://github.com/Illumina/paragraph) and python 3.10.13. The JSON files used to describe variants for GGTyper were converted to VCF files containing reference and alternate sequences with margin 150 bp and to JSON files containing graph descriptions using custom python scripts.
For simulated samples, read depth across chromosomes 20 and 21 was calculated using

*samtools coverage*[5]. For the Polaris data, we used the target sequencing coverage of 30x as input. In the simulations, we used BAM files containing all reads simulated from the two chromosomes as input. The Polaris BAM files we used contained only variant regions with a margin of 10 000 bp. We used the complete human reference genome GRCh38.p13 as reference.

**VCF input.** Using a single VCF file to describe 20 variants, we genotyped 199 Polaris samples using *multigrmpy*.

**Graph input.** Alternatively it is possible to supply graph structures as JSON files to Paragraph. We called *grmpy* separately for each variant graph. All other parameters remained identical.

**Filtering.** When filtering of variant calls was applied, we used a threshold of $GQ = 20$.

###### 2.10.2 BayesTyper

We use BayesTyper [6] in v1.5 (https://github.com/bioinformatics-centre/BayesTyper, static build) and KMC [7] v3.2.2 (https://github.com/refresh-bio/KMC). We prepared a VCF file containing reference and alternate allele sequences for all variants as for Paragraph, but additionally left-aligned and normalized the VCF file using *bcftools norm* as specified on the BayesTyper GitHub page.
We followed the instructions on GitHub to create $k$-mer bloom filters from BAM files for use with BayesTyper before calling *bayesTyper cluster* and *bayesTyper genotype*.
For the real data, we used the BayesTyper GRCh38 bundle in version 1.3 for the canon and decoy sequences. In our simulations, we had to swap the canon file for a fasta file containing only the sequences of chromosomes 20 and 21 in order to allow accurate estimation of expected $k$-mer counts even when simulating from an incomplete reference genome.
The default filtering did not result in usable genotype calls for our 20 complex SVs in the Polaris data. Due to that we removed all filters from the VCF file using *bayesTyperTools filter* as described on GitHub.

**Filtering.** On simulated data, we left the default filters in place and additionally used the same quality threshold of $GQ = 20$ as for Paragraph. When testing BayesTyper with default filters on 10 samples from the Polaris data for 20 real variants, all variant calls with at least one variant allele were removed during filtering. We therefore removed all default filters and only used $GQ = 20$ during filtering. For reasons unknown to us, removing all filters using *bayesTyperTools filter* removed NA_variant_7 from the VCF file. Therefore, no results exist for this variant.

### 3 Variants

#### 3.1 Simulated variants

##### 3.1.1 Simulation details and variant origin

SVs of all canonical (Fig. S12) and six different complex types (Fig. S13) were simulated. csvA, csvB, csvC and csvD are based on real variants associated with Mendelian disorders [8]. csvI is a variation on csvC and csvT is a made up translocation with additional complications around the breakpoints.
Variant positions are uniformly distributed on chromosomes 20 and 21, excluding the centromere region of chromosome 21 and any N-containing sections in the hg38 reference sequence. Variants were simulated without overlap, with separating margins of at least 1 kbp.
The sizes of the individual segments are drawn randomly with minima and maxima determined by the variant templates used for simulation, which can be found on GitHub. Segments are generally at least 50 bp in size and the largest simulated segments have size 10 kbp. DNA segments within a variant structure have different maximum sizes, emulating the variants that the structures are based on.

##### 3.1.2 Structures

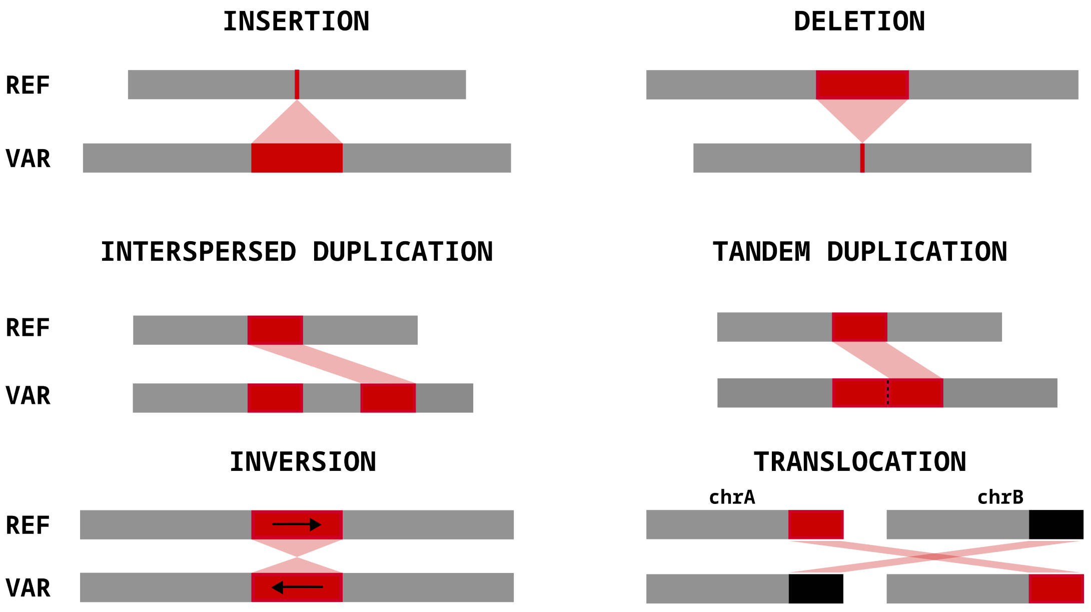

Figure S12: Canonical SV types.

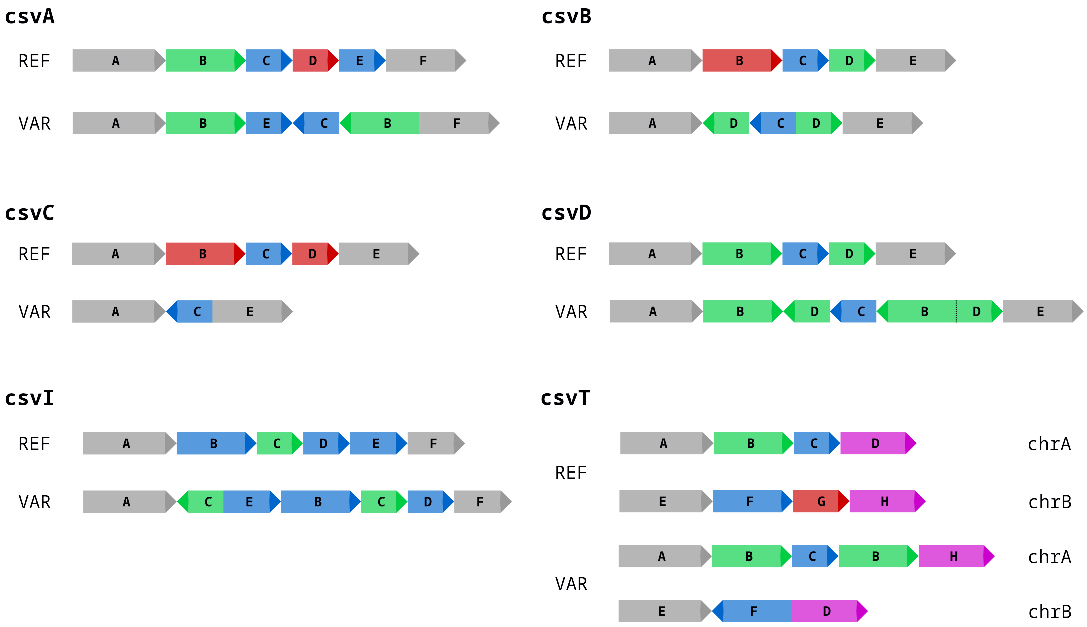

Figure S13: Simulated complex SV structures.

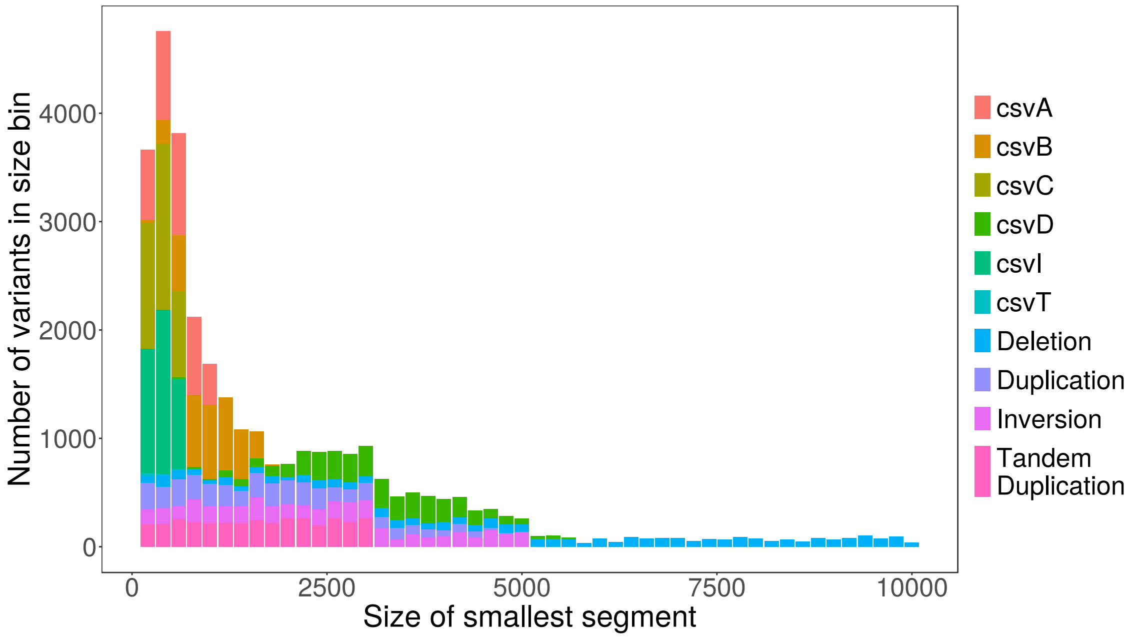

Figure S14: Histogram of smallest segment size per variant across variant types.

#### 3.2 Real variants

##### 3.2.1 Origins and locations

Table 1: Complex SVs used for evaluation on real data.

| ID | Name | Location | Structure | Origin |
|---|---|---|---|---|
| V1 | NA_Variant_1 | chr1:205209503-205209666 | Inv-Del | [9] |
| V2 | NA_Variant_2 | chr2:10685912-10687069 | Inv-Dup | [9] |
| V3 | NA_Variant_3 | chr3:95746924-95752156 | Del-DispDup | [9] |
| V4 | NA_Variant_4 | chr5:79750309-79756108 | DispInvDup-DispDup | [9] |
| V5 | NA_Variant_5 | chr8:99145584-99146021 | Inv-Dup | [9] |
| V6 | NA_Variant_6 | chr10:125501849-125512653 | DispInvDup-DispDup-DispDup | [9] |
| V7 | NA_Variant_7 | chr17:5691379-5694179 | Del-PartInv-Del | [9] |
| V8 | svVariant_1 | chr1:187495696-187497598 | Inv-Del | [9] |
| V9 | svVariant_2 | chr2:124294179-124295663 | DispInvDup | [9] |
| V10 | svVariant_3 | chr3:80013716-80016128 | Del-DispInvDup | [9] |
| V11 | svVariant_4 | chr5:28932544-28934740 | Del-DispInvDup | [9] |
| V12 | svVariant_5 | chr9:62806329-62822795 | Del-Inv-Del | [9] |
| V13 | svVariant_6 | chr9:86539642-86541032 | Del-Inv | [9] |
| V14 | svVariant_7 | chr10:57497185-57498224 | Del-Inv | [9] |
| V15 | svVariant_8 | chr2:61473618-61476323 | DispInvDup | [9] |
| V16 | svVariant_9 | chr9:105054356-105055067 | DispInvDup | [9] |
| V17 | svVariant_10 | chr12:77990098-77994497 | DispInvDup | [9] |
| V18 | Spink14 | chr5:148173476-148175216 | Del-Inv-Del | [10] |
| V19 | ShortInversion | chr3:131989409-131994540 | Del-Inv-Del | [10] |
| V20 | SVA-E Insertion | chr2:201281936-201284717 | Del | [11] |

*Del*: deletion, *Dup*: duplication, *Inv*: inversion, *DispDup*: dispersed duplication, *DispInvDup*: dispersed inverted duplication, *PartInv*: partial inversion.

Schematic structures shown in Fig. S15. All coordinates refer to the human reference chromosome GRCh38.p13. SVA-E is technically a canonical deletion, but with the additional difficulty of possible copies of the same sequence on other chromosomes. Most variants were taken from a paper by Lin et al. [9]. Among the variants they listed in their supplementary materials, we selected those that were indicated as complex, with breakpoints recognizable in short read data. Subsequent manual verification and – if necessary – adjustment of breakpoint positions and structures (see Fig. S15) followed. Due to the time-consuming task of manual verification and structure determination, we stopped after gathering 20 variants in total.

##### 3.2.2 Structures

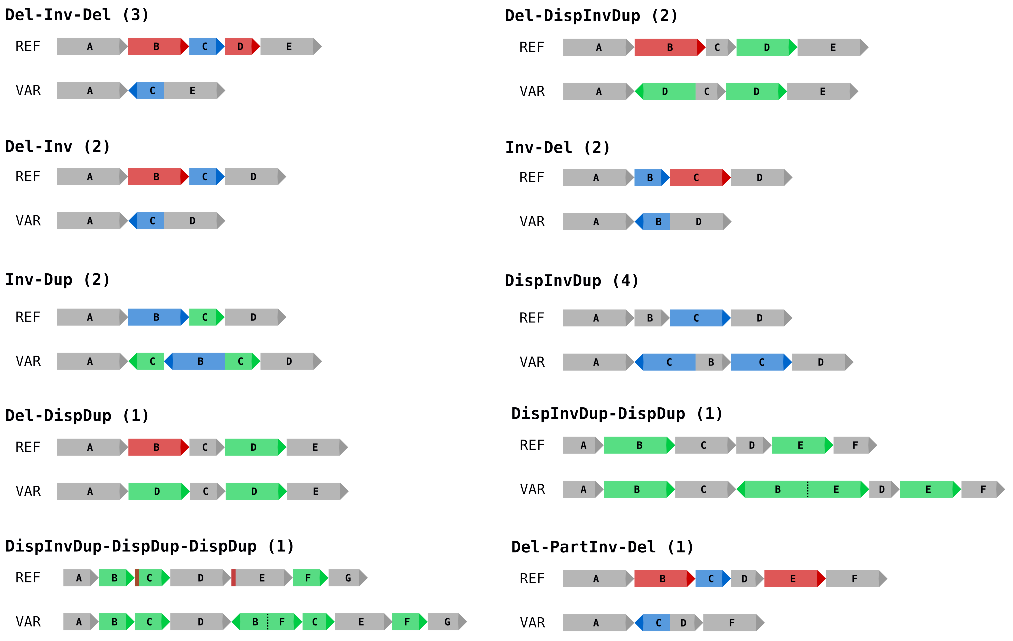

Figure S15: Rough structures of real complex SVs listed in Tab. 1. Small deletions (mainly at novel junctions) may be missing in the figure.
